# Identifiability investigation of within-host models of acute virus infection

**DOI:** 10.1101/2024.05.09.593464

**Authors:** Yuganthi R. Liyanage, Nora Heitzman-Breen, Necibe Tuncer, Stanca M. Ciupe

## Abstract

Uncertainty in parameter estimates from fitting within-host models to empirical data limits the model’s ability to uncover mechanisms of infection, disease progression, and to guide pharmaceutical interventions. Understanding the effect of model structure and data availability on model predictions is important for informing model development and experimental design. To address sources of uncertainty in parameter estimation, we use four mathematical models of influenza A infection with increased degrees of biological realism. We test the ability of each model to reveal its parameters in the presence of unlimited data by performing structural identifiability analyses. We then refine the results by predicting practical identifiability of parameters under daily influenza A virus titers alone or together with daily adaptive immune cell data. Using these approaches, we present insight into the sources of uncertainty in parameter estimation and provide guidelines for the types of model assumptions, optimal experimental design, and biological information needed for improved predictions.

**Author summary:** Within-host models of virus infections fitted to data have improved our understanding of mechanisms of infection, disease progression, and allowed us to provide guidelines for pharmaceutical interventions. Given their predictive power, it is essential that we properly uncover and address uncertainty in model predictions and parameter estimation. Here, we focus on the effect of model structure and data availability on our ability to uncover unknown parameters. To address these questions, we use four mathematical models of influenza A infection with increased degrees of biological realism. We test the ability of each model to reveal its parameters in the presence of unlimited data by performing structural identifiability analysis. We then refine the results by predicting practical identifiability of parameters under daily influenza A virus titers alone or together with daily adaptive immune cell data. Using these approaches, we present insights into the sources of uncertainty in parameter estimations and provide guidelines for the types of model assumptions, optimal experimental design, and biological information needed for improved predictions.

## Introduction

The study of host-virus interactions using dynamical models (within-host models) has improved our understanding of the mechanistic interactions that govern chronic [1–4] and acute [5–12] viral infections. Regardless of the virus considered, the most basic within-host model has a general structure that includes the interaction between the cells susceptible to the virus, the cells infected by the virus, and the virus at short (acute) and long (chronic) time-scales. The emergence of unexpected dynamics in the virus data, new information about the virus’ life-cycle, data describing host immunity to the infection, or a combination of some or all of the above, may require addition of complexity into the within-host modeling process (see [13] and [14] for a review).

Data fitting techniques for simple or complex within-host models use (normalized) least-squares approaches, in which the Euclidean distance between the data and the mathematical model is minimized with respect to the unknown parameters. The first step in the parameter estimation algorithm is to provide an initial guess for each parameter based on prior biological knowledge. When prior knowledge is unknown, the user makes the assumption that any positive parameter guess is acceptable. Then an optimization search algorithm is employed until a termination criterion is reached. For many within-host mathematical models and corresponding data sets, the optimization is ill-posed due to the structure of the model and/or the frequency of the data [15]. As a result, some parameters may be difficult or impossible to quantify. To determine whether the uncertainty in parameter estimations is due to the model or the data, both structural and practical identifiability questions need to be addressed.

Structural identifiability investigates the ability of a model to reveal its unknown parameters from noise-free infinite amount of data [16–19]. When non-structural identifiability of parameters occurs, it is important to find the source of non-identifiability, such as correlation between model parameters. This allows the user to propose additional assumptions needed to make the model structurally identifiable. Only after the structural identifiability of the unknown parameters is guaranteed, one can conduct data fitting schemes to estimate parameter values.

Practical identifiability investigates the ability of a model to reveal unknown structurally identifiable parameters under scarce and noisy (among subjects) data [16, 19–24, 24–28]. As with the structural identifiability, it is important to identify whether the practical identifiability issues are due to model structure. Additionally, it is important to determine whether increased data frequency, availability of data measurements for more than one model variable, and/or relaxing restrictions imposed on the unknown parameters can improve practical identifiability issues.

To address these important considerations in model validation, one needs to compare a set of models for the same virus infection system and the same empirical data. Here, we accomplish that by investigating four previously developed models of influenza A virus (IAV) infection in mice [29]. The first three models, all validated with the same virus titer data set, are ranging from the basic within-host model to models with increased complexity through the addition of non-linear terms and/or the inclusion of additional variables for the host cell populations infected by the influenza virus. The fourth model is the most complex, due to the addition of both non-linear terms and variables for the host immune system. This results in a large number of unknown parameters. To compensate for the added complexity, this model is validated with two data sets: the same virus titer data and an additional immune cell population data.

The goal of this study is to determine how model complexity and data availability induce uncertainty in parameter estimates. Using the proposed models as proof of concept, we aim to provide a framework for model validation, from structural to practical identifiability, that can be generalized to other models of virus infections.

## Materials and methods

### Mathematical models

We consider four within-host models of acute infections used to describe influenza A virus infection in mice [6]. The flow charts of the four models are presented in Fig. **??**.

<u>Model 1</u> is the classical target-cell limitation model of viral infection which considers the interaction between target cells, infected cells and virus, as follows [6, 29]. Target cells, *T*, interact with the virus, *V* at rate *β* to become infected cells *I*. Infected cells die at per capita rate *δ* and produce virus at rate *π*. Virus is cleared at rate *c*. <u>Model 1</u> is described by the system of ordinary differential equations (ODE) Eq. 1 below,

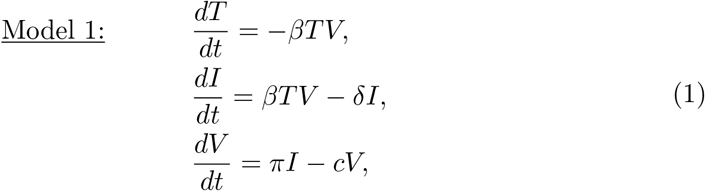

with initial conditions *T* (0) = *T*_0_, *I*(0) = *V*_0_, and *V* (0) = 0.

Experimental data has shown that, following peak expansion, virus decays in a biphasic manner. To capture the dynamics of viral decay, a modified death rate has been considered. It assumes that the rate of infected cell clearance increases as the density of infected cells decreases, as described by *δ/*(*K*_*δ*_ + *I*) [29], where *δ* is the maximum per capita death rate and *K*_*δ*_ is infected cell population where death rate is half-maximal. This leads to the modified target-cell limitation <u>Model 2</u> given by the ODE system Eq. 2 below,

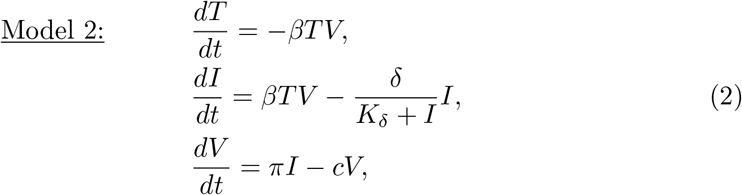

with initial conditions *T* (0) = *T*_0_, *I*(0) = *I*_0_, and *V* (0) = 0.

It was observed experimentally that, following influenza A virus exposure, there is a delay between infection of target cells and viral production by infected cells [30]. The delay was accounted for by assuming that, upon infection, cells enter an eclipse phase *I*_1_, where cells are infected but do not produce virus. They become productively infected *I*_2_ after 1*/k* days [5], where 1*/k* is the average time spent in eclipse phase. This leads to the target-cell limitation model with eclipse phase <u>Model 3</u> given by the ODE system Eq. 3 below,

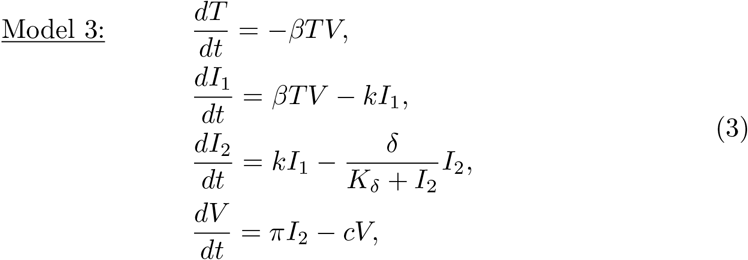

with initial conditions *T* (0) = *T*_0_, *I*_1_(0) = 0, *I*_2_(0) = *I*_0_, and *V* (0) = 0.

The first three models do not explicitly account for any immune responses, but indirectly assume infected cell loss at non-specific rate *δ* (or *δ/*(*K*_*δ*_ + *I*_2_)) and viral clearance at non-specific rate *c*. The observed biphasic viral decay captured by <u>Models 2-3</u> given by Eq. 2 - Eq. 3, however, have the additional feature that the timing of the second phase viral decay coincides with the development of adaptive immune cells in the form of CD8^+^ T cells, which are responsible for killing infected cells and resolving the infection [6]. To account for adaptive immunity (especially in the presence of immune cell data), an additional variable *E* is considered. It only accounts for the effector CD8^+^ T cell population (and ignores the memory CD8^+^ T cell population), as follows. In the absence of infection, a baseline of influenza A virus-specific effector CD8^+^ T cells are present, *E*(0) = *E*_0_. Infection results in recruitment of additional effector CD8^+^ T cells at a rate proportional to the productively infected cells *I*_2_. This is modeled in a density dependent manner at rate *λ/*(*K*_*E*_ + *E*), where *λ* is the maximum influx and *K*_*E*_ is the effector CD8^+^ T cell population where the influx is half-maximal. Effector CD8^+^ T cells proliferate in the presence of infection. This is modeled by a delayed term *ηI*_2_(*t* − *τ*_*I*_)*E*, which assumes that expansion occurs following interaction between effector CD8^+^ T cells and cells that became productively infected *τ*_*I*_ days ago. To account for effector CD8^+^ T cells function, the model assumes that effector CD8^+^ T cells kill infected cells in a density dependent manner modeled by the term *δ*_*E*_*/*(*K*_*δ*_ + *I*_2_), where *δ*_*E*_ is the maximum per capita killing rate and *K*_*δ*_ is the *I*_2_ concentration where the killing is half-maximal. A non-specific infected cell killing rate *δ* is still considered. The resulting delay differential equations (DDE) immune model is described by the DDE system Eq. 4 below,

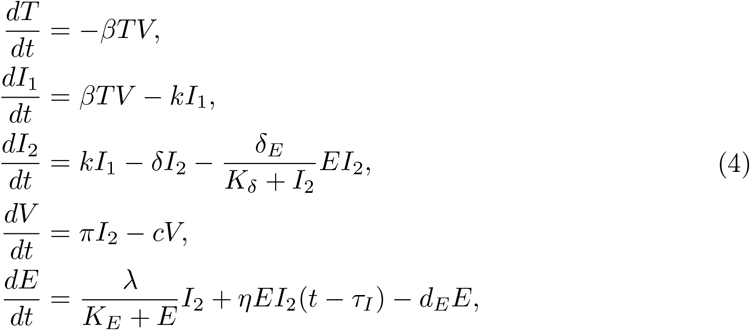

with initial conditions *T* (0) = *T*_0_, *I*_1_(0) = *I*_0_, *V* (0) = 0, *E*(0) = *E*_0_, and *I*_2_(*t*) = 0 for −*τ*_*I*_ ≤ *t* ≤ 0.

To unify the results by investigating uncertainty in parameter estimates when fitting ODE virus dynamics systems to data, we first approximate the DDE system given by Eq. 4 with an ODE system, as follows [31]. For a delay of *τ*_*I*_ days, we incorporate *n* dummy variables which all span *τ*_*I*_ */n* days in the variable *I*_2_’s dynamics. Briefly, we let *y*_*i*_ be the productively infected cell populations at times 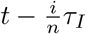 days post infection, for *i* = 1, …, *n* and consider the following ODE system for dummy variables *y*_*i*_(*t*),

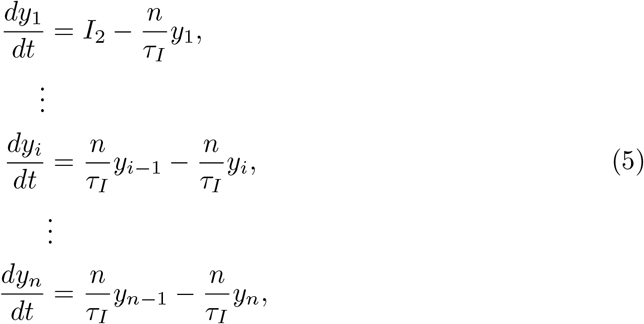

with *y*_*i*_(0) = 0 for *i* = 1, …, *n*. Then the delayed productively infected cell population is given by

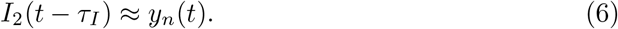

Without loss of generality, we assume *n* = 3. The corresponding immune <u>Model 4</u> is given by the ODE system Eq. 7 below,

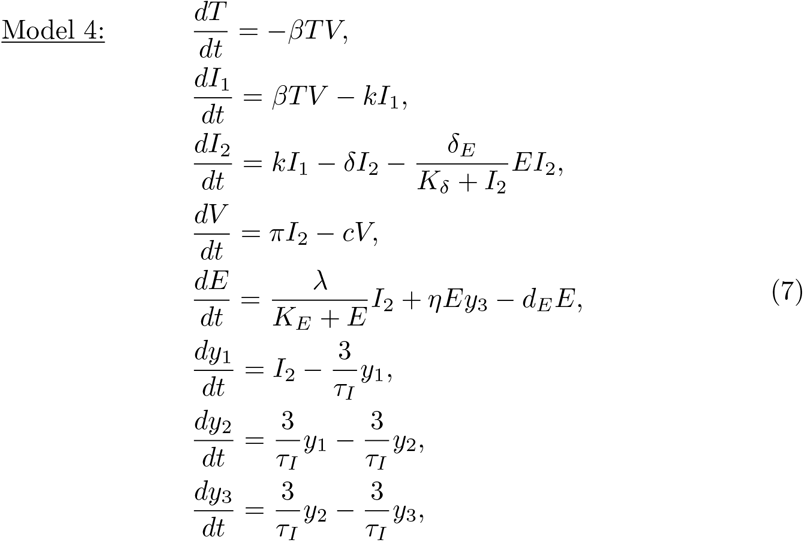

with initial conditions *T* (0) = *T*_0_, *I*_1_(0) = *I*_0_, *I*_2_(0) = 0, *V* (0) = 0, *E*(0) = *E*_0_, and *y*_*i*_(0) = 0 for *i* = 1, 2, 3.

### Structural identifiability theory

To study the structural identifiability of the <u>Models 1-4</u>, we rewrite them in the following general form:

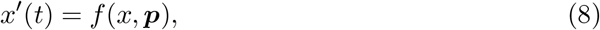

and the observations as

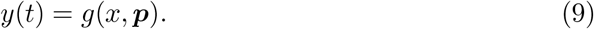

Here, *x* denotes the state variables, ***p*** is the parameter vector and *y* is the output (given by the empirical data), also called the observations. The generic model given by Eq. 8 is assumed structurally identifiable if the parameter vector ***p*** can be determined uniquely from the given observations *y*(*t*), assumed to be unlimited. Otherwise, it is said to be unidentifiable. The formal definition of structural identifiability is provided below.

#### Definition 1.

*Let* ***p*** *and* 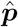 *be two distinct parameter vectors. Model Eq. 8 is said to be globally (uniquely) structurally identifiable if*

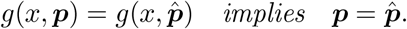

#### Definition 2.

*Model Eq. 8 is said to be locally structurally locally identifiable if for any* ***p*** *within an open neighborhood of*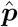 *in the parameter space*,

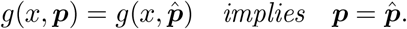

Various methods have been proposed for analyzing the structural identifiability of ODE models [16–18]. In this study, we employ differential algebra approach. It performs the elimination of unobserved state variables, resulting in equations expressed as functions of model parameters and observed state variables. These are referred to as the input-output equations, and are differential-algebraic polynomials consisting of the outputs, *y*(*t*), with model parameters, ***p*** as coefficients. The formal definition of structural identifiability within differential algebra approach for model Eq. 8 is provided below.

#### Definition 3.

*Let c*(***p***) *denote the coefficients of the input-output equation corresponding to model Eq. 8*. *We say that model Eq. 8 is structurally identifiable from unlimited observations y*(*t*) *if and only if*

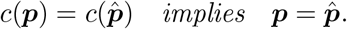

Studying structural identifiability of ODE models using the differential algebra methods can be accomplished using several platforms and available open-source software. Here we presents three such platforms: the Differential Algebra for Identifiability of System (DAISY) [32], the Identifiable **Comb**inations web application (COMBOS) [33], and the StructuralIdentifiability.jl in JULIA [34].

There are many similarities among the three methods. All of them offer insights into the structural identifiability status of each parameter by categorizing them into locally identifiable, globally identifiable, or non-identifiable. They employ a differential elimination method to calculate input-output equations of the considered system, and test the one-to-one map between the coefficients of the input-output equations and model parameters. COMBOS and StructuralIdentifiability.jl package in JULIA are superior to DAISY, as they provides globally identifiable parameter correlations, in an otherwise non-identifiable system. Even though DAISY does not print parameter correlations, they can be derived using the coefficients of the input-output equations and algebraic manipulations in software such as MATHEMATICA. Of the three software, COMBOS does not print the input-output equations, making for a faster (yet more opaque) platform. Previous studies have shown that COMBOS works best for small to medium-size models and is not assured for models with large parameter vectors [33, 35, 36]. While highly similar, it is up to the user to determine which software is best suited for studying the identifiability of the models considered.

### Data fitting methods

#### Empirical data

We use previously published longitudinal influenza A infectious virus titer and CD8^+^ T cell data in mice from *Smith, et al*. [6]. Adult mice were inoculated intranasally with 75 TCID_50_ of mouse adapted influenza A/Puerto Rico/8/34 (H1N1) (PR8) virus.

Total infectious virus (log 10 TCID_50_ per lung) was measured for ten mice each day. Nine days after inoculation, infectious virus was no longer detectable in any of the mice. Therefore, we only consider infectious virus titer data from the first nine days post inoculation in our analyses. We let 𝔼 (*V*_*data*_(*i*)) be the mean infectious virus titer data at day *i* = {1, …, 9} and 𝕍**ar**(*V*_*data*_(*j*)) be the infectious virus titer variance at days *i* = {1, …, 9} among the ten mice.

Moreover, total effector CD8^+^ T cells (cells per lung) were measured daily for five mice. Since influenza A-specific effector CD8^+^ T cells were detectable for all twelve days of the study, we consider effector CD8^+^ T cells data from the first twelve days post inoculation in our analyses. We let 𝔼 (*E*_*data*_(*j*)) be the population mean CD8^+^ T cell data (per lung) at day *j* = {1, …, 12} and 𝕍**ar**(*E*_*data*_(*j*)) be the CD8^+^ T cell data variance at days *j* = {1, …, 12} among the five mice.

#### Model parameters and initial conditions

For all models, we assume known initial conditions *T* (0) = 10^7^ cells/ml, *I*(0) = 75 cells/ml, and *V* (0) = 0 virus per ml as in [6]. For <u>Models 3-4</u>, we additionally assume that *I*_2_(0) = 0. For <u>Model 4</u>, we assume *E*(0) = *E*_0_ is unknown, therefore added *E*_0_ to the parameter vector to be estimated from the data. Lastly, we assume all parameters are unknown. When parameters are either very large or very small, we estimate their value on natural logarithmic scale. In particular, we estimate **p**_**1**_= {ln(*β*), *δ, π, c*} for <u>Model 1</u>, **p**_**2**_= {ln(*β*), ln(*δ*), ln(*K*_*δ*_), *π, c*} for <u>Model 2</u>, **p**_**3**_= {ln(*β*), ln(*δ*), ln(*K*_*δ*_), *π, c, k*} for <u>Model 3</u> and **p**_**4**_= {ln(*β*), *δ*, ln(*K*_*δ*_), *π, c, k, δ*_*E*_, ln(*η*), ln(*λ*), *d*_*E*_, ln(*K*_*E*_), *τ*_*I*_, ln(*E*_0_)} for <u>Model 4</u>.

### Data fitting algorithm

To estimate parameters **p**_**1**_-**p**_**3**_, we fit the predicted viral load log_10_ 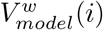 given by <u>Models 1-3</u> to the longitudinal mean (among the ten mice) infectious virus (log_10_ TCID_50_ per lung) 𝔼 (*V*_*data*_(*i*)), knowing that the variance in the data at day *i* is 𝕍**ar**(*V*_*data*_(*i*)), for *i* = {1…9} days. We assume that the data satisfies the following statistical model [21, 37]

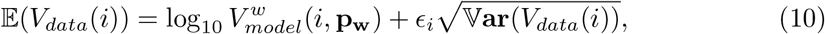

where, 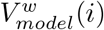 is the predicted virus trajectory given by <u>Model w</u> at day *i* post infection, *i* = {1, …, 9} days, and **p**_**1**_= {ln(*β*), *δ, π, c*}, **p**_**2**_= {ln(*β*), ln(*δ*), ln(*K*_*δ*_), *π, c*} and **p**_**3**_= {ln(*β*), ln(*δ*), ln(*K*_*δ*_), *π, c, k*}, and *ϵ*_*i*_ are independent and identically distributed with mean zero and standard deviation *σ*. Given the statistical model Eq. 10, we assume that the measured data, 𝔼 (*V*_*data*_(*i*)), follows a normal distribution with a mean equal to the model prediction 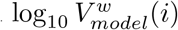 and with variance equal to *σ*^2^ 𝕍**ar**(*V*_*data*_(*i*)). Moreover, the availability of measurements from several animals that vary with time, allows us to account for changing variance in data over time, 𝕍**ar**(*V*_*data*_(*i*)). Therefore, we consider the following functional (weighted residual sum of squares), to estimate the model parameters,

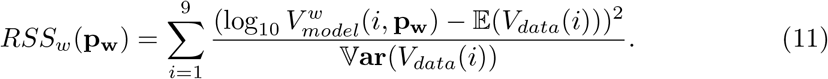

Then the parameters for <u>Models 1-3</u> are estimated by minimizing the weighted least-squares given by,

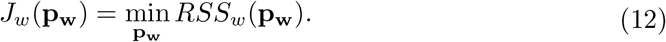

Moreover, to estimate parameters **p**_**4**_, we fit both the predicted viral load 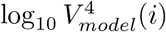 given by <u>Model 4</u> to the longitudinal mean (among ten mice) infectious virus 𝔼 (*V*_*data*_(*i*)), knowing that the variance in the data at day *i* is 𝕍**ar**(*V*_*data*_(*i*)), for *i* = {1…9} days and the predicted effector cell population 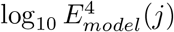 given by <u>Model 4</u> to the longitudinal mean (among five mice) CD8^+^ T cell data 𝔼 (*E*_*data*_(*j*)), knowing that the variance in the data at day *j* is 𝕍**ar**(*E*_*data*_(*j*)), for *j* = {1…12} days. We assume that the data satisfy the following the statistical model [21, 37]

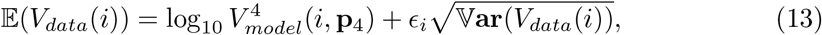

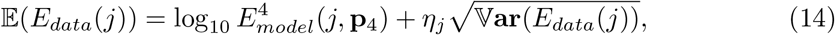

where, 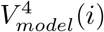 is the predicted virus trajectory given by <u>Model 4</u> at day *i* post infection, *i* = {1, …, 9} days, 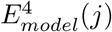 is the predicted CD8^+^ T cell population given by <u>Model 4</u> at day *j* post infection, *j* = {1, …, 12} days, and **p**_**4**_= {ln(*β*), *δ*, ln(*K*_*δ*_), *π, c, k, δ*_*E*_, ln(*η*), ln(*λ*), *d*_*E*_, ln(*K*_*E*_), *τ*_*I*_, ln(*E*_0_)}. Here, *ϵ*_*i*_ and *η*_*j*_ are independent and identically distributed with mean zero and standard deviations *σ*_*V*_ and *σ*_*E*_, respectively. As before, the measured data, 𝔼 (*E*_*data*_(*j*)) follows a normal distribution whose mean is the model prediction 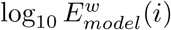 and with variance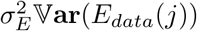. Consider the following functional (weighted residual sum of squares),

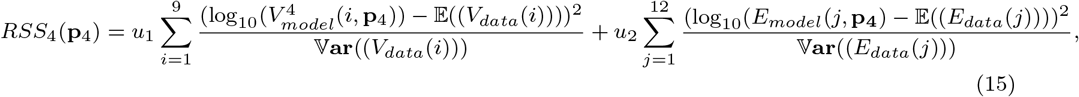

Then the parameters for <u>Model 4</u> are estimated by minimizing the weighted least-squares given by,

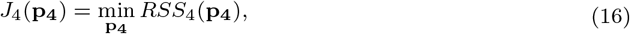

Note that we weighted the virus and effector cells contributions, with weights *u*_1_ = 1 and *u*_2_ = max_*j*_ 𝕍**ar**(*E*_*data*_(*j*))*/* max_*i*_ 𝕍**ar**(*V*_*data*_(*i*)).

We minimize the least-square functionals *RSS*_*w*_ using the *fmincon* function in MATLAB with ranges for parameters **p**_**w**_ given in Table 1.

**Table 1.** Upper and lower bounds for parameters estimated by fitting <u>Models 1-3</u> to influenza virus and by fitting <u>Model 4</u> to both influenza virus and effector CD8^+^ T cell data from infected mice.

| Parameter | Model 1 | Model 2 | Model 3 | Model 4 |
| --- | --- | --- | --- | --- |
| $\beta \times 10^{-5}$ | 0.1 – 10 | 0.1 – 10 | 0.1 – 10 | 0.1 – 10 |
| $\delta$ | 0-25 | $10^2$-$10^8$ | $10^2$-$10^8$ | 0-15 |
| $K_\delta$ | - | $10^3$-$10^7$ | $10^3$-$10^7$ | $10^1$-$10^5$ |
| $\pi$ | $0$-$10^2$ | $0$-$10^2$ | $0$-$10^2$ | $0$-$10^2$ |
| $c$ | 0-25 | 0-25 | 0-25 | 0-25 |
| $k$ | - | - | 4-6 | 4-6 |
| $\delta_E$ | - | - | - | 0-25 |
| $\eta \times 10^{-7}$ | - | - | - | $10^{-2}$ – 100 |
| $\lambda \times 10^3$ | - | - | - | $10^{-2}$-$10^2$ |
| $d_E$ | - | - | - | 0-25 |
| $K_E \times 10^6$ | - | - | - | $0.1$ – $10^3$ |
| $\tau_I$ | - | - | - | 0-10 |
| $E_0 \times 10^3$ | - | - | - | $0.1$ – 10 |

#### Model selection

To compare <u>Models 1-4</u>, we calculate the corrected Akaike Information Criteria (AICc), given below,

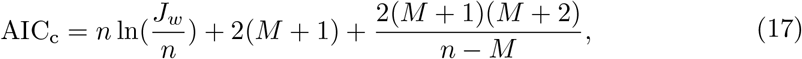

where *n* is the number of data points used to estimate parameters **p**_**w**_ and *M* is the number of estimated parameters. In <u>Models 1-3</u>, *n* = 9 and *M* = 4, 5 and 6, respectively. In Model 4, *n* = 21 and *M* = 13.

#### Model prediction confidence interval

To quantify the uncertainty associated with predicted solutions of each model, we perform parametric bootstrapping. It is a simulation-based method which assumes that data comes from a known distribution with unknown parameters. For <u>Models 1-3</u>, we assume that the predicted viral population for best parameter estimates, 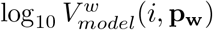, is the mean of the data’s normal distribution and *σ*^2^ 𝕍**ar**(*V*_*data*_(*i*)) is its variance (see Eq. 10). Then *σ* can be approximated as follows,

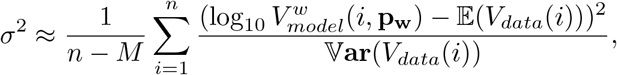

(see Banks et al. for a full derivation [37]). Here, *n* = 9 is the number of viral samples and *M* is the number of parameters (*M* = 4, *M* = 5 and *M* = 6 for <u>Models 1-3</u>, respectively). To find a confidence region in our model predictions, we generate 1000 simulated datasets using the distribution space given by Eq. 10, and fit Models 1-3 to each data sets.

Similarly, for <u>Model 4</u>, assuming that viral data and effector cell data come from distributions with means 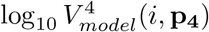 and 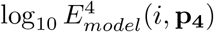 (the predicted variables for best parameter fits) and that 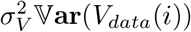 and 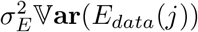 are the variances then,

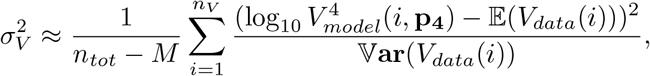

and

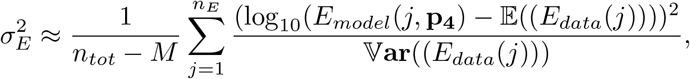

as before. Here *n*_*V*_ = 9 is the number of viral samples, *n*_*E*_ = 12 is the number of CD8^+^ T cell samples, *n*_*tot*_ = *n*_*V*_ + *n*_*E*_ = 21 is the total data samples and *M* = 13 is the number of parameters fitted.

### Practical identifiability theory

While structural identifiability investigates whether parameters can be uniquely determined from a model given unlimited data in the absence of measurement error or noise, practical identifiability determines whether parameters can be accurately identified in real-world scenarios, where observed discrete and variable among subject data is contaminated with measurement errors. We and others have employed several methods to study practical identifiability of within-host and infectious disease models, such as *Monte Carlo Simulations* [19–21], the *Fisher Information Matrix* (FIM) or *Correlation Matrix* [16, 22–24], *and Bayesian Methods* [25]. In this study we use the *Profile Likelihood Method* [26, 38] *described in detail below*.

Consider that the vector of parameters **p** is partitioned into **p***=(r*, **s**), where *r* represents the parameter whose practical identifiability we are investigating and **s** represents the vector of remaining parameters. The profile likelihood of *r* is given by,

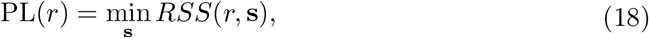

where *RSS* is the objective functional used for data fitting (in our case Eq. 11 for <u>Models 1-3</u> and Eq. 15 for <u>Model 4</u>). In other words, PL(*r*) finds the minimum of the objective functional *RSS*(*r, s*) for an array of fixed *r* values over the space of the remaining parameters **s**. The shape of PL(*r*) informs the identifiability of *r*, with a u-shaped PL(*r*) that exceeds a threshold (corresponding to a chosen confidence level) indicating practical identifiability of *r* and a flat PL(*r*) indicating non-practical identifiability of *r*.

We estimate PL(*r*) over a mesh of equally spaced values of *r*, centered at the best-fit estimate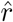, with the number of mesh points chosen to have enough data to generate a confidence interval for 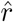, as follows. If we consider a model with parameters **p** unknown and a model with parameters **s** unknown, we obtain two nested models that differ by parameter *r*. It has been shown that the likelihood ratio of the nested models converges to a *χ*^2^ distribution with one degree of freedom, *df* = 1 (see *Corrollary 2* in [38] for more detail). This helps us define the Δ-level likelihood-based confidence interval for parameter *r* to be

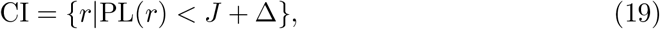

where Δ is the percentile of the *χ*^2^ distribution with *df* = 1, and *J* is the weighted least-squares functional at the best parameter estimate [38]. This can be summarized, as follows.

#### Definition 4.

*Let CI* = {*r*|*PL*(*r*) *< J* + Δ} *be the likelihood-based confidence interval for parameter r*.

1. *If CI ⊂* [*r*_1_, *r*_2_] *where r*_1_ *and r*_2_ *are finite, then parameter r is practically identifiable*.
2. *If either r*_1_ *or r*_2_ *is infinite, then parameter r is not practically identifiable*.

*A model is practically identifiable if all parameters are practically identifiable*.

For <u>Models 1-3</u>, we generated the profile likelihoods PL(*r*) for each parameter *r* ∈ {**p**_**w**_} for **w** = {1, 2, 3} by fitting the functional *RSS*_*w*_(**p**_**w**_) given by Eq. 11 and <u>Model **w**</u> to mean population viral load data. We obtained best estimates for the remaining parameters **s** over a mesh of equally spaced, known *r* values. Similarly, for <u>Model 4</u>, we generated the profile likelihood PL(*r*) for unknown parameters *r* ∈ {**p**_**4**_} by fitting functional *RSS*_4_(**p**_**4**_) given by Eq. 15 simultaneously to the mean population viral load and the mean population effector cell data. We obtained best estimates for the remaining parameters **s** over a mesh of equally spaced, known *r* values.

Additionally, to further explore the relationship between data availability and practical identifiability, we generated profile likelihoods for parameters **p**_**w**_ and <u>Model **w**</u> using simulated, noise-free, high frequency data sets. In particular, we assumed that the virus load data was collected every fourteen minutes (for a total of 899 evenly spaced points) for <u>Models 1-3</u> and that both the virus load data and the effector cell data were collected every fourteen minutes (for a total of 899 evenly spaced points of each data type) for <u>Model 4</u>. We fitted each model to this high frequency data and generated profile likelihoods of the resulting parameter values. In all of our models we chose Δ to be the 90-th percentile of the *χ*^2^-distribution, 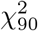 [38]. This guaranteed us a 90% confidence interval in the estimated parameter *r*.

## Results

To investigate the effect of model structure, model complexity, and data availability on the uncertainty in model prediction and parameter estimation, four ordinary differential equation models of influenza A virus infection are being considered. <u>Model 1</u>, given by Eq. 1, assumes simple interactions between target epithelial cells, infected epithelial cells, and the influenza A virus using mass action and linear terms of interaction. <u>Model 2</u>, given by Eq. 2, expands on model Eq. 1 by assuming a density dependent infected cell death rate. <u>Model 3</u>, given by Eq. 3, expands on model Eq. 2 by adding an equation for exposed (but not productively infected) epithelial cells. Lastly, <u>Model 4</u>, given by Eq. 7, adds several variables and terms of interaction for the immune cell development and function, modeled by linear, mass-action, and density dependent terms of interaction (see “Mathematical models” in **Materials and methods**). We investigated the structural identifiability of each model, performed data fitting to either virus titer data alone or to combined viral titer and effector CD8^+^ T cell data, and investigated practical identifiablity of the estimated parameters.

### Structural identifiability results

To determine whether the considered models can reveal their parameters, we examine the structural identifiability of <u>Models 1-3</u>, given by Eqs. 1-3, under unlimited observations of viral load and the structural identifiability of <u>Model 4</u>, given by Eq. 7, under unlimited combined observations of viral load and effector CD8^+^ T cell concentrations. We used the differential algebra software DAISY (see “Structural identifiability theory” in **Materials and methods** for details).

### DAISY-based structural identifiability results for <u>Model 1</u>

We assume all <u>Model 1</u> parameters ***p*** = {*β, δ, π, c*} are unknown and that we have unlimited empirical observations of the viral load, *y*(*t*) = *V* (*t*). Using DAISY [32], we obtain the following input-output equation in variable *V* and model parameters ***p***,

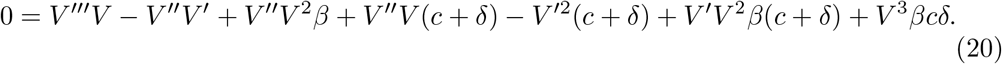

By Definition 3 (see “Structural identifiability theory” in **Materials and methods**), we need to examine whether another set of parameters, 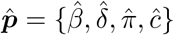 can produce the same empirical observation *V* (*t*), making the map from the parameter space ***p*** to the coefficients of input-output equation Eq. 20 one-to-one. If we set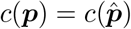, we obtain

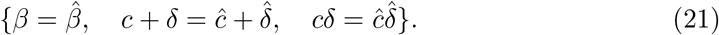

Solving Eq. 21 results in the following two sets of solutions;

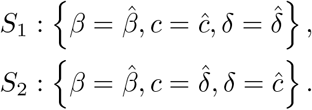

Hence, only the infection rate *β* is globally structurally identifiable, while the infected cells killing rate *δ*, and the virus clearance rate *c* are locally identifiable. Lastly, the virus production rate *π* does not appear in the input-output equation Eq. 20. Therefore, it is not structurally identifiable. We summarize the results for <u>Model 1</u> below (see Table 2).

**Table 2.** Structural identifiability results for <u>Models 1-4</u> using three software: DAISY, StructuralIdentifiability.jl package in JULIA and COMBOS.

| Model | Observed States | DAISY | JULIA | COMBOS |
| --- | --- | --- | --- | --- |
| <u>Model 1</u><br>unknown initial conditions | $V(t)$ | $\beta$<br>globally identifiable<br>$\{c, \delta\}$<br>locally identifiable<br>Correlations:<br>$c + \delta = \hat{c} + \hat{\delta}$,<br>$c\delta = \hat{c}\hat{\delta}$ | $\beta$<br>globally identifiable<br>$\{c, \delta\}$<br>locally identifiable<br>$\pi$<br>nonidentifiable<br>Correlations:<br>$c + \delta = \hat{c} + \hat{\delta}$,<br>$c\delta = \hat{c}\hat{\delta}$ | $\beta$<br>globally identifiable<br>$\{c, \delta\}$<br>locally identifiable with 2 solutions |
| <u>Model 2</u><br>unknown initial conditions | $V(t)$ | $\{\beta, c\}$<br>globally identifiable<br>Correlations:<br>$\delta\pi = \hat{\delta}\hat{\pi}$, $\pi K_\delta = \hat{\pi}\hat{K}_\delta$ | $\{\beta, c\}$<br>globally identifiable<br>$\{\pi, K_\delta, \delta\}$<br>nonidentifiable<br>Correlations:<br>$\delta\pi = \hat{\delta}\hat{\pi}$, $\pi K_\delta = \hat{\pi}\hat{K}_\delta$ | $\{\beta, c, K_\delta\pi, \delta\pi\}$<br>globally identifiable |
| <u>Model 3</u><br>unknown initial conditions | $V(t)$ | $\{\beta, c, k\}$<br>globally identifiable<br>Correlations:<br>$\frac{\delta}{K_\delta} = \frac{\hat{\delta}}{\hat{K}_\delta}$, $\pi K_\delta = \hat{\pi}\hat{K}_\delta$ | $\{\beta, c, k\}$<br>globally identifiable<br>$\{\pi, K_\delta, \delta\}$<br>nonidentifiable<br>Correlations:<br>$\delta\pi = \hat{\delta}\hat{\pi}$, $\pi K_\delta = \hat{\pi}\hat{K}_\delta$ | $\{\beta, c, K_\delta\pi, \delta\pi\}$<br>globally identifiable |
| <u>Model 4</u><br>unknown initial conditions | $V(t), E(t)$ | $\{\beta, c, d_E, \delta, k, K_E, \tau_I\}$<br>globally identifiable<br>Correlations:<br>$\delta_E\eta = \hat{\delta}_E\hat{\eta}$,<br>$K_\delta\eta = \hat{K}_\delta\hat{\eta}$,<br>$\frac{\lambda}{\eta} = \frac{\hat{\lambda}}{\hat{\eta}}$, $\frac{\pi}{\eta} = \frac{\hat{\pi}}{\hat{\eta}}$ | $\{\beta, c, d_E, \delta, k, K_E, \tau_I\}$<br>globally identifiable<br>$\{\delta_E, K_\delta, \pi, \lambda, \eta\}$<br>nonidentifiable<br>Correlations:<br>$K_\delta\pi = \hat{K}_\delta\hat{\pi}$,<br>$K_\delta\lambda = \hat{K}_\delta\hat{\lambda}$,<br>$\eta K_\delta = \hat{\eta}\hat{K}_\delta$,<br>$\delta_E\pi = \hat{\delta}_E\hat{\pi}$ | identifiability unknown |
| **All models**<br>known initial conditions except for $E_0$ | $V(t)$<br>$V(t), E(t)$ | <u>Models 1-3</u><br>globally identifiable<br><u>Model 4</u><br>identifiability unknown | <u>Models 1-4</u><br>globally identifiable | <u>Models 1-4</u><br>identifiability unknown |

#### Proposition 1.

*<u>Model 1</u> given by Eq. 1 is not structured to identify all of its parameters from unlimited viral load observations, V* (*t*). *More precisely, parameter β is globally structurally identifiable, parameters c and δ are locally structurally identifiable and parameter π not structurally identifiable. Moreover, <u>Model 1</u> is globally structural identifiable under known initial conditions*.

### DAISY-based structural identifiability results for <u>Model 2</u>

We assume all parameters ***p*** = {*β, δ, K*_*δ*_, *π, c*} for <u>Model 2</u>, given by Eq. 2, are unknown and that we have unlimited empirical observations of the viral load, *y*(*t*) = *V* (*t*). Using DAISY, we obtain the following input-output equation,

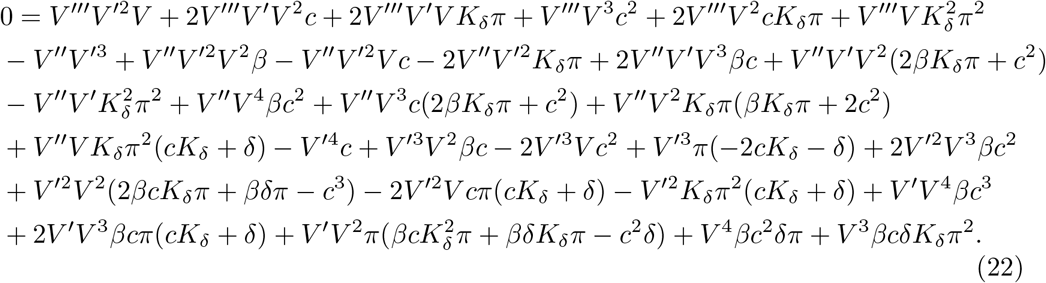

As before, we examine whether another set of parameters, 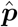, can produce the same empirical observation *V* (*t*), making the map from the parameter space ***p*** to the coefficients of input-output equation Eq. 22 one-to-one. If we set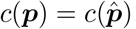, we obtain

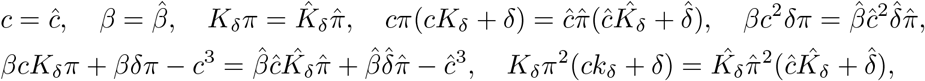

with solutions

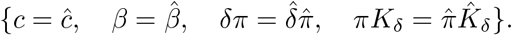

Hence, <u>Model 2</u> is not structurally identifiable. In particular, infection rate *β*, viral clearance rate *c*, and the products *δπ, K*_*δ*_*π* (but not the individual parameters *δ, π* and *K*_*δ*_) are globally identifiable. Since the correlations *δπ* and *K*_*δ*_*π* are known, fixing one of these parameters can make model Eq. 2 identifiable. We summarize the results for <u>Model 2</u> below (see Table 2).

#### Proposition 2.

*<u>Model 2</u> given by Eq. 2 is not structured to identify all of its parameters from unlimited viral load observations, V* (*t*). *More precisely, parameters β and c are globally structurally identifiable. Moreover the parameter products δπ and K*_*δ*_*π are globally identifiable. Since the correlations are known, fixing δ, π or K*_*δ*_ *makes the <u>Model 2</u> globally structurally identifiable from unlimited observations V* (*t*). *Moreover, <u>Model 2</u> is globally structural identifiable under known initial conditions*.

### DAISY-based structural identifiability results for Model 3

We assume all parameters ***p*** = {*β, δ, k, δ*_*E*_*K*_*δ*_, *π, c*} for Model 3, given by Eq. 3, are unknown and that we have unlimited empirical observations of the viral load, *y*(*t*) = *V* (*t*). Using DAISY, we can derive the input-output equations (they are too messy and will not be presented here). As before, we examine whether another set of parameters, 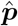 can produce the same empirical observation *V* (*t*), making the map from parameter space ***p*** to coefficients of input-output equation (not shown) one-to-one. If we set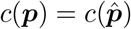, we obtain

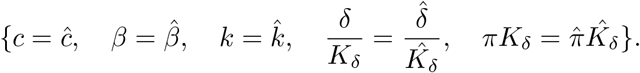

Hence, <u>Model 3</u> is not structurally identifiable. In particular, the infection rate *β*, the eclipse parameter *k*, the viral clearance rate *c*, the ratio *δ/K*_*δ*_ and the product *K*_*δ*_*π* (but not the individual parameters *δ, π* and *K*_*δ*_) are globally identifiable. Since the correlations are known, fixing one of these parameters makes the model Eq. 3 identifiable. We summarize the results for <u>Model 3</u> below (see Table 2).

#### Proposition 3.

*<u>Model 3</u> given by Eq. 3 is not structured to identify all of its parameters from unlimited viral load observations, V* (*t*). *More precisely, parameters β, k and c are globally structurally identifiable. Moreover the parameters ratio δ/K*_*δ*_ *and product K*_*δ*_*π are globally identifiable. Since the correlations are known, fixing the parameter δ, π or K*_*δ*_ *makes the <u>Model 3</u> globally structurally identifiable from unlimited observations V* (*t*). *Moreover, <u>Model 3</u> is globally structural identifiable under known initial conditions*.

### DAISY-based structural identifiability results for <u>Model 4</u>

To study the structural identifiability of <u>Model 4</u> (given by Eq. 7) we assume that all parameters, ***p*** = {*β, δ, k, δ*_*E*_*K*_*δ*_, *π, c λ, η, d*_*E*_, *τ*_*I*_, *E*_0_}, are unknown and that we have unlimited empirical observations for the viral load *y*_1_(*t*) = *V* (*t*) and the effector cell data *y*_2_(*t*) = *E*(*t*). Using DAISY, we can obtain input-output equation (they are messy and will not be presented here). As before, we examine whether another set of parameters,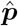 can produce the same empirical observation *V* (*t*) and *E*(*t*), making the map from the parameter space ***p*** to the coefficients of input-output equation (not shown) one-to-one. If we set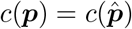, we obtain

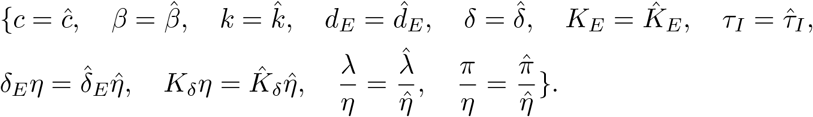

Hence, <u>Model 4</u> is not structurally identifiable. In particular, the infection rate *β*, the eclipse parameter *k*, the viral clearance rate *c*, the effector cells death rate *d*_*E*_, the generic killing rate *δ*, the half-maximal level *K*_*E*_, the delay *τ*_*I*_, the ratios *λ/η* and *π/η* and the products *d*_*E*_*η* and *K*_*δ*_*η* (but not the individual parameters *δ*_*E*_, *η, K*_*δ*_, *π, λ*) are globally identifiable. If the parameter *η* is fixed, then the model Eq. 7 becomes identifiable. We summarize the results for <u>Model 4</u> below (see Table 2).

#### Proposition 4.

*<u>Model 4</u> given by Eq. 7 is not structured to identify all of its parameters from unlimited viral load and effector cell observations, V* (*t*) *and E*(*t*). *More precisely, parameters β, k, c, d*_*E*_, *δ, K*_*E*_, *and τ*_*I*_ *are globally structurally identifiable. Moreover the parameter ratios λ/η and π/η and parameter products d*_*E*_*η and K*_*δ*_*η are globally identifiable. If the parameter η is fixed, then <u>Model 4</u> becomes globally structurally identifiable from unlimited observations V* (*t*) *and E*(*t*).

We do not know (from DAISY) whether knowing initial conditions guarantees global stability of <u>Model 4</u> (see Table 2).

### Comparison among structural identifiability software

Studying structural identifiability of ODE models can be achieved using software other than DAISY (see “Structural identifiability theory” in **Materials and methods**). To determine how these methods compare, results from three platforms DAISY, COMBOS [33], and StructuralIdentifiability.jl in JULIA [34], for <u>Models 1-4</u> are presented side by side in Table 2.

We find that all three software uncover the same identifiability results for <u>Models 1-3</u>. On the other hand, DAISY and StructuralIdentifiability.jl in JULIA uncover the same identifiability results (while COMBOS cannot find results) for <u>Model 4</u> under unknown initial conditions. Even though <u>Model 3</u> and <u>Model 4</u> employ different interpretations of the parameter correlations among platforms, simple algebraic manipulations show that the obtained correlations are equivalent. Given the similarity in the results among <u>Models 1-3</u>, it is up to the user to decide which of the three software is best suited for their analysis. Similarly, given the similarity in the results among DAISY and StructuralIdentifiability.jl in JULIA for <u>Model 4</u> with unknown initial conditions, it is up to the user to decide which of the two software is best suited for its analysis. However, only StructuralIdentifiability.jl in JULIA can be used to determine the structural identifiability of <u>Model 4</u> with unknown *E*_0_ and known other initial conditions. Hence, for larger systems with non-linear terms of interactions this method should be employed.

## Data fitting results

We fitted <u>Models 1-3</u> to previously published longitudinal influenza A infectious virus load and we fitted <u>Model 4</u> to both longitudinal influenza A infectious virus load and longitudinal CD8^+^ T cell data in infected mice [6], using a normalized least-square optimization algorithm (see “Data fitting methods” in **Materials and methods**). The results from fitting *V* (*t*) given by <u>Models 1-3</u> to viral load data are shown in Fig. 2**A-C** and the best parameter fits are given in Table 3. The results from fitting both *V* (*t*) and *E*(*t*) given by <u>Model 4</u> to viral load and effector cell data are shown in Fig. 2**D** and the best parameter fits are given in Table 3. Model selection, using the corrected Akaike Information Criteria (*AIC*_*c*_), predicts that <u>Model 4</u> best describes the data (see Table 3). To quantify the uncertainty associated with predicted solutions of each model, we find a 95% confidence regions in our model predictions (see “Model prediction confidence interval” in **Materials and methods**), illustrated by shaded grey areas in Fig. 2**A-C** for <u>Models 1-3</u> and by grey and blue shaded region in Fig. 2**D**, for <u>Model 4</u>. We see large error regions in virus population predictions for all models during the decay phase (grey regions in Fig. 2**A-D**). Moreover, <u>Models 2-4</u> better capture the virus population expansion phase compared to <u>Model 1</u> (grey regions in Figs. 2**B-D** versus grey region in Fig. 2**A**). Lastly, the largest error in CD8^+^ T cell prediction in <u>Model 4</u> occurs in the second week of infection (blue region in Fig. 2**D**).

**Fig 1.**
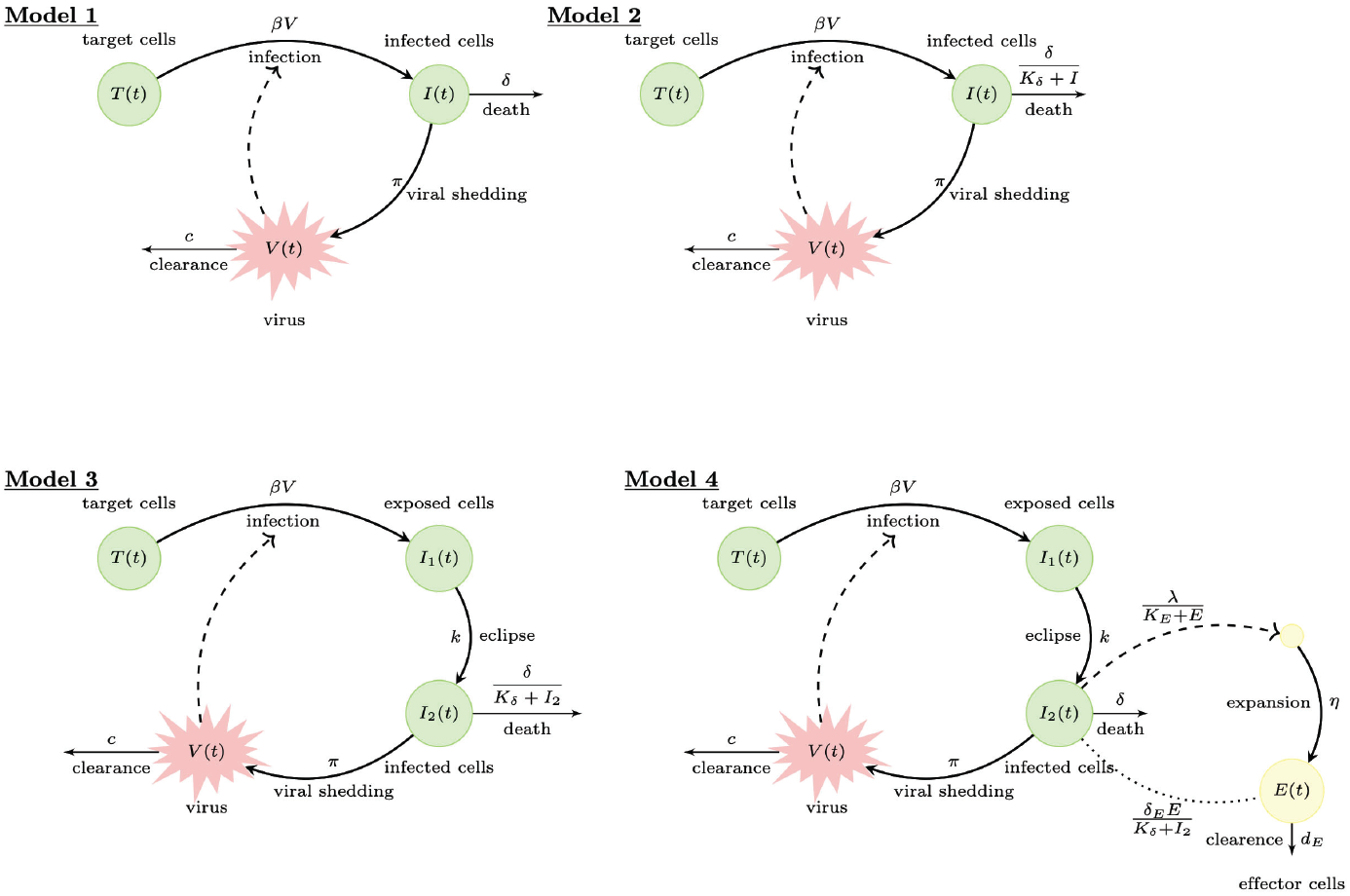
Flow charts for Models 1-4.

**Table 3.** Parameter estimates found by fitting <u>Model 1</u>, <u>Model 2</u>, and <u>Model 3</u> to virus load data and <u>Model 4</u> to virus load and effector CD8^+^ T cell data from mice infected with influenza A virus using *fmincon* routine in MATLAB.

| Parameter | Model 1 | Model 2 | Model 3 | Model 4 |
| --- | --- | --- | --- | --- |
| $\beta \times 10^{-5}$ | 0.48 | 6.88 | 6.20 | 8.39 |
| $\delta$ | 1.47 | $1.59 \times 10^6$ | $1.72 \times 10^6$ | 0.196 |
| $K_\delta$ | - | $4.69 \times 10^4$ | $2.37 \times 10^5$ | $5.49 \times 10^3$ |
| $\pi$ | 1.18 | 0.86 | 1.69 | 0.69 |
| $c$ | 1.48 | 6.49 | 12.34 | 6.19 |
| $k$ | - | - | 4.82 | 5.02 |
| $\delta_E$ | - | - | - | 14.20 |
| $\eta \times 10^{-7}$ | - | - | - | 3.59 |
| $\lambda \times 10^3$ | - | - | - | 1.31 |
| $d_E$ | - | - | - | 0.20 |
| $K_E \times 10^6$ | - | - | - | 9.68 |
| $\tau_I$ | - | - | - | 1.69 |
| $E_0 \times 10^3$ | - | - | - | 1.21 |
| $J_w$ | 21.12 | 2.09 | 2.39 | 2.14 |
| AIC<sub>c</sub> | 37.67 | 40.88 | 114.1 | 36.78 |

**Fig 2.**
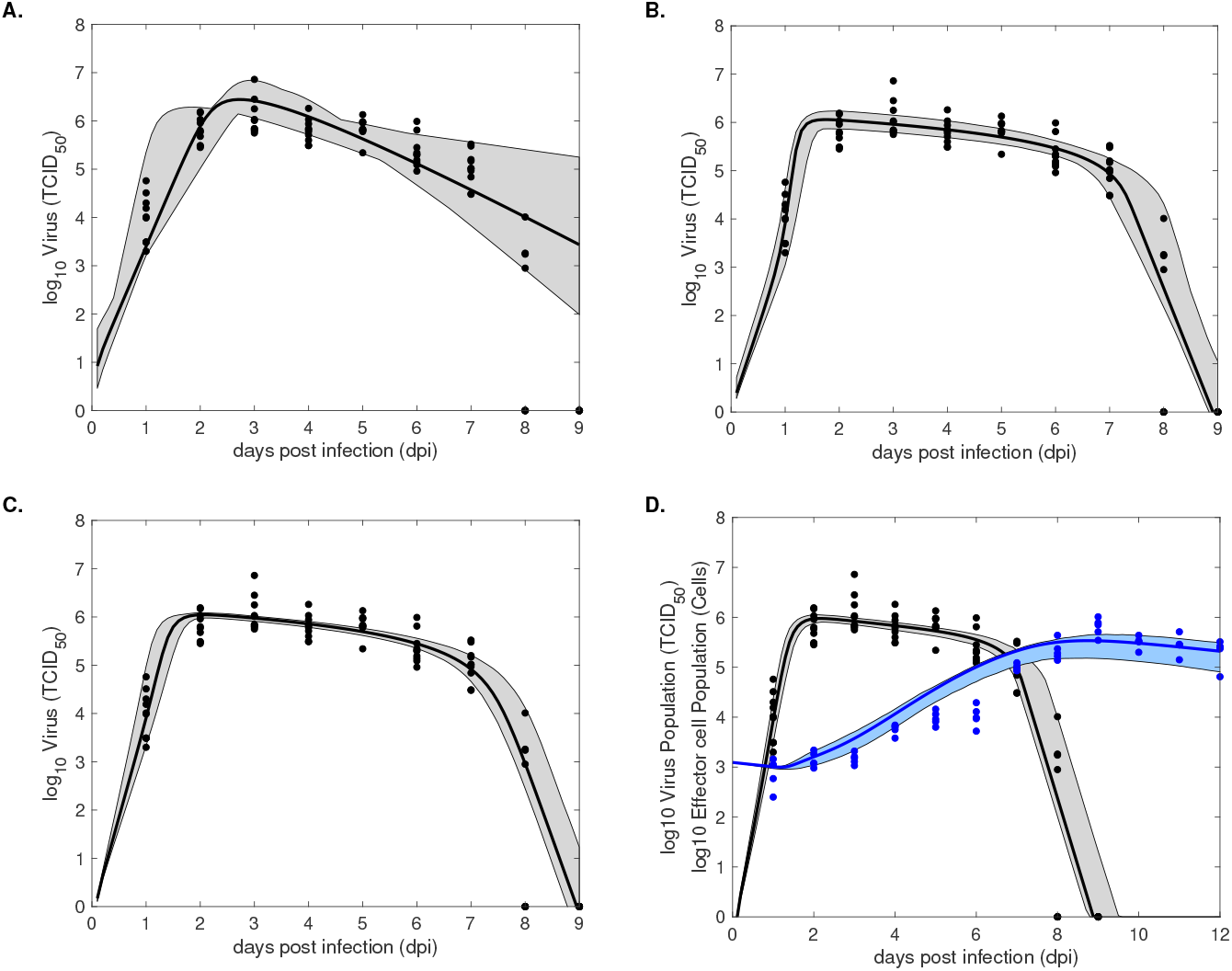
Model predictions (solid lines) and 95% model confidence regions (dashed areas) obtained by fitting *V* (*t*) (black lines) given by **A:** <u>Model 1</u>, **B:** <u>Model 2</u>, **C:** <u>Model 3</u> to virus load data (black circles) and by fitting *V* (*t*) (black line) and *E*(*t*) (blue line) given by **D:** <u>Model 4</u> to o virus load data (black circles) and CD8^+^ T cell data (blue circles). Model parameters are given in Table 3.

### Practical identifiability results

Since we determined that <u>Models 1-4</u> are structurally identifiable under known initial conditions and unlimited data, we were allowed to search for best estimates for all models’ parameters. The resulting fitting routine may still be ill-posed, given that the data consisted of discrete (daily) data sets that varied among the infected mice, rather than the unlimited data required by the structural identifiability theory. Therefore, we performed practical identifiability for <u>Models 1-4</u> under the discrete subject data in [29] (see “Practical identifiability theory” in **Materials and methods**). We used the *Profile Likelihood* practical identifiability method [24, 26–28], which has the advantage of not only determining whether a parameter is practically identifiable, but also of determining a 90% confidence interval for the parameter space where practical identifiability occurs (see “Practical identifiability theory” in **Materials and methods**).

When using the empirical population mean virus titer data in [6] for Model 1, we found that *β* is practically identifiable with 90% confidence interval *CI ⊂* [1.64 *×* 10^−6^, 2.94 *×* 10^−5^] and *π* is practically identifiable with 90% confidence interval *CI ⊂* [0.31, 3.18], respectively. On the other hand, both *δ* and *c* are not practically identifiable, with identical 90% confidence intervals *CI ⊂* [0.87, ∞] (see Fig. 3**A**). Adding high frequency data and rerunning the profile likelihood analysis resulted in practical identifiability of all four parameters, consistent with the structural identifiability results for <u>Model 1</u> (see Fig. 3**B**).

**Fig 3.**
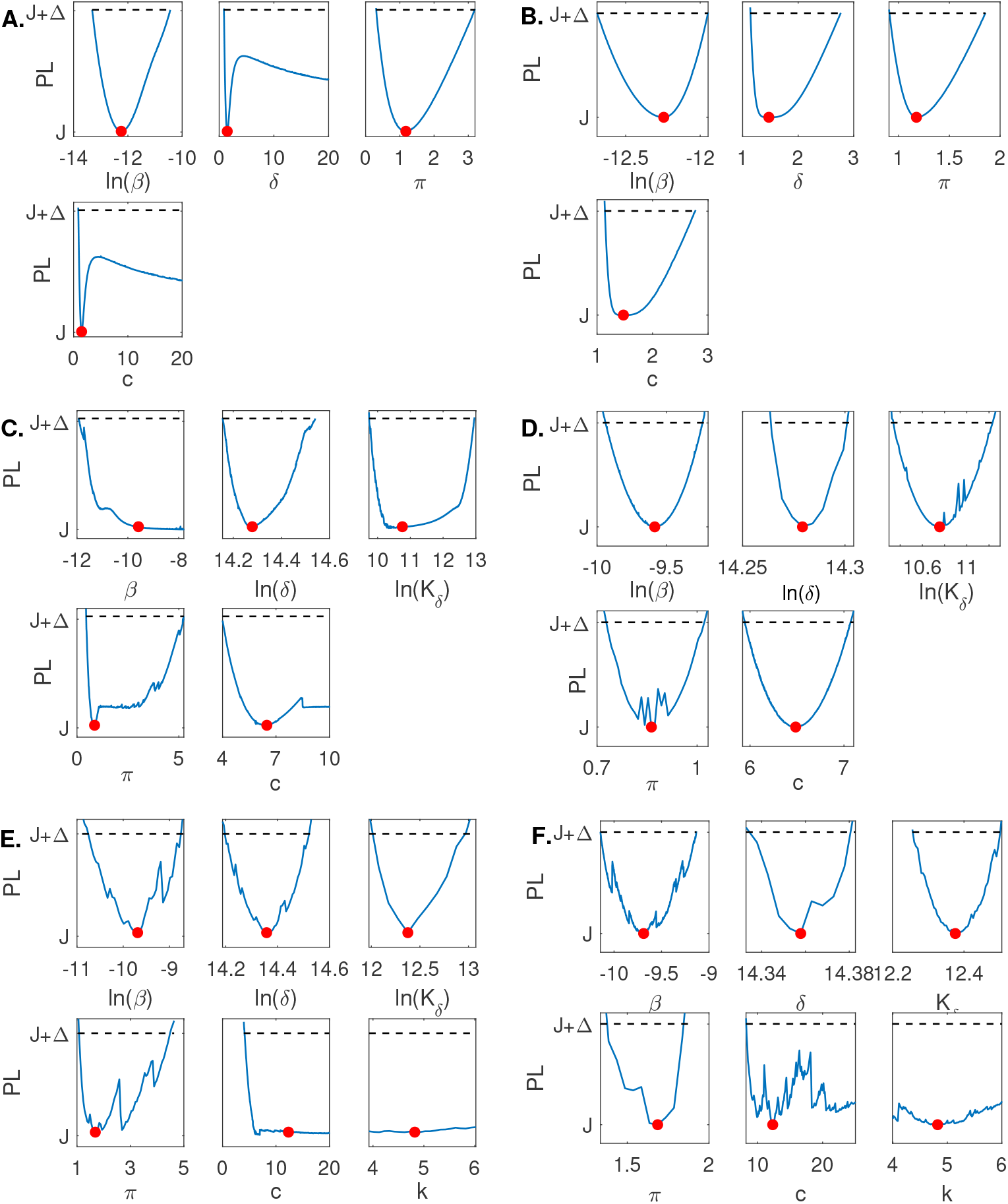
Profile likelihood curves generated using empirical data for **A:** <u>Model 1</u>, **C:** <u>Model 2</u>, **E:** <u>Model 3</u>; and profile likelihood curves generated using high frequency simulated data for **B:** <u>Model 1</u>, **D:** <u>Model 2</u>, **F:** <u>Model 3</u>. The red circles indicate best parameter estimates given in Table 3 and the dashed lines represent a threshold equivalent to 90% confidence level in the parameter estimate.

Similar to <u>Model 1</u>, when using the empirical population mean virus titer data in [6] for <u>Model 2</u>, we found that *δ, K*_*δ*_, and *π* are practically identifiable with 90% confidence intervals *CI ⊂* [1.41 *×* 10^6^, 2.06 *×* 10^6^], *CI ⊂* [1.74 *×* 10^4^, 4.19 *×* 10^5^], and *CI ⊂* [0.44, 5.24], respectively. On the other hand, both *β* and *c* are not practically identifiable, with 90% confidence intervals *CI ⊂* [6.62 *×* 10^−6^, ∞] and *CI ⊂* [3.98, ∞], respectively (see Fig. 3**C**). Adding high frequency data and rerunning the profile likelihood analysis resulted in practical identifiability of all five parameters, consistent with the structural identifiability results for <u>Model 2</u> (see Fig. 3**D**).

For <u>Model 3</u>, when using the empirical population mean virus titer data in [6], we found that *β, δ, K*_*δ*_, and *π* are practically identifiable with 90% confidence intervals *CI ⊂* [2.17 *×* 10^−5^, 1.60 *×* 10^−4^], *CI ⊂* [1.47 *×* 10^6^, 2.03 *×* 10^6^], *CI ⊂* [1.59 *×* 10^5^, 4.32 *×* 10^5^], and *CI ⊂* [1.07, 4.49], respectively. On the other hand, both *c* and *k* are not practically identifiable, with 90% confidence intervals *CI ⊂* [3.98, ∞] and *CI ⊂* [∞, ∞], respectively (see Fig. 3**E**). Adding high frequency data and rerunning the profile likelihood analysis did not result in practical identifiability of all five parameters (see Fig. 3**F**). However, if we additionally relaxed constraints on parameters *c* and *k* to range in the [0, 1000] and [0, 50] intervals, compared to the constraints chosen in Table 1, we observed practical identifiability of all five parameters, consistent with the structural identifiability results for Model 3 (see Fig.5**B**).

For <u>Model 4</u>, when using the discrete empirical population mean virus titer data and the empirical population mean effector cell data in [6] simultaneously, we found that *π* and *E*_0_ are practically identifiable with 90% confidence intervals *CI ⊂* [0.29, 2.49] and *CI ⊂* [66, 5.41 *×* 10^−4^]. Parameters *k, λ*, and *K*_*E*_ are not practically identifiable with the same 90% confidence interval *CI ⊂* [−∞, ∞]. Parameters *β, δ*_*E*_, *K*_*δ*_, *η*, and *c* are also not practically identifiable with 90% confidence intervals *CI ⊂* [8.00 *×* 10^−6^, ∞], *CI ⊂* [1.10, ∞], *CI ⊂* [42.98, ∞], *CI ⊂* [5.37 *×* 10^−8^, ∞], and *CI ⊂* [3.89, ∞], respectively. Lastly, parameters *δ, d*_*E*_, and *τ* are not practically identifiable on the positive domain with an undefined lower bound (ULB) for the 90% confidence interval and a finite upper bound. In particular, *CI ⊂* [*ULB*, 0.63], *CI ⊂* [*ULB*, 3.20], *CI ⊂* [*ULB*, 6.99], for *δ, d*_*E*_ and *τ*_*I*_, respectively (see Fig. 4**A**). Adding high frequency data and rerunning the profile likelihood analysis resulted in practical identifiability of all thirteen parameters, consistent with the structural identifiability results for <u>Model 4</u> (see Fig. 4**B**).

**Fig 4.**
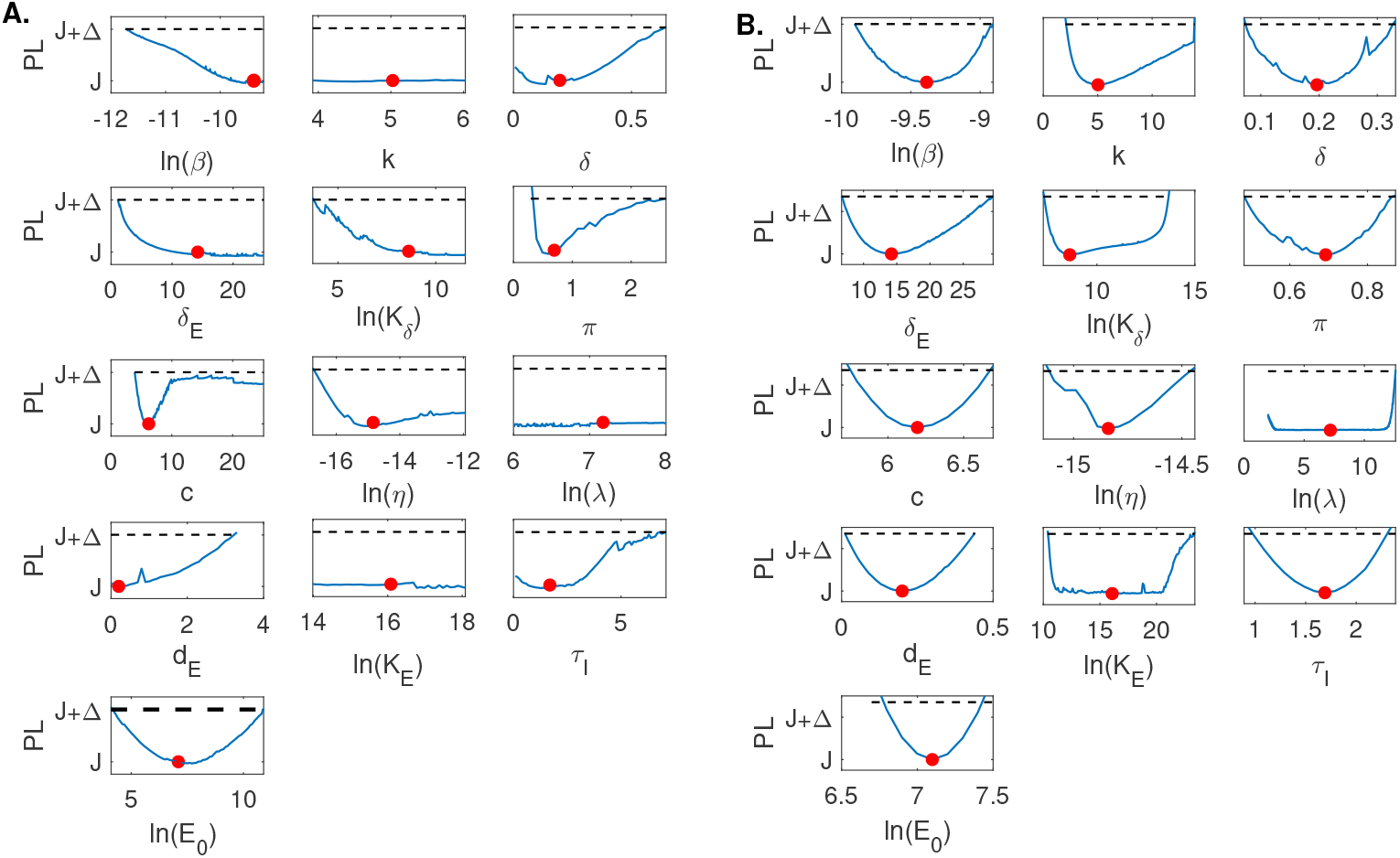
Profile likelihood curves generated using empirical data for **A:** <u>Model 4</u>, and profile likelihood curves generated using high frequency simulated data for **B:** <u>Model 4</u>. The red circles indicate best parameter estimates given in Table 3 and the dashed lines represent a threshold equivalent to 90% confidence level in the parameter estimate.

## Discussion

In this study, we investigated the conditions needed to ensure model identifiability in four models of influenza A virus dynamics in infected mice. To apply the same methodology and software, all considered models were either modeled by systems of ordinary differential equations (<u>Models 1-3</u> given by equations Eq. 1-3) or approximated by a system of ordinary differential equations (<u>Model 4</u> given by Eq. 7). The considered models differ in the number of equations (corresponding to the number of variables) from three in <u>Models 1-2</u>, to four in <u>Model 3</u> to eight in <u>Model 4</u>. Consequently, the number of unknown parameters is a maximum of four in <u>Model 1</u>, a maximum of five in Model 2, a maximum of six in <u>Model 3</u> and a maximum of thirteen in <u>Model 4</u>. Lastly, the terms of interaction include only mass-action and linear terms for <u>Model 1</u> and mass-action, linear terms, and density dependence terms for <u>Models 2-4</u>.

We found that the increased complexity, needed to capture biological realism, comes with a cost. It resulted in increased uncertainty in parameter estimates not only when discrete and noisy virus and immune cell empirical data is used for validation but also when we assumed (hypothetically) that unlimited data is available. This means that data fitting should not be conducted until it is established that parameters can be revealed from unlimited data under the considered model structure. In other words, the first step in the model validation is determining whether all unknown parameters are structurally identifiable (see Fig.6).

When it comes to investigating the structural identifiability of systems of ordinary differential equations several software platforms are available. Here, we compared results from three of them; DAISY [32], COMBOS [33] and StructuralIdentifiability.jl in JULIA [34]. For <u>Model 1-3</u> and unlimited virus titer data, we found the same classification for the structurally identifiable parameters and the same (or equivalent) correlations between the non-structurally identifiable parameters, regardless of the software used (Table 2). For the more complex <u>Model 4</u> and unlimited virus titer and effector CD8^+^ T cell data, however, only StructuralIdentifiability.jl in JULIA finds that the model is structurally identifiable under known (with the exception of initial effector population, *E*_0_) initial conditions (Table 2). When initial conditions are unknown, we find identical classification for structurally identifiable parameters and equivalent correlations between the non-structurally identifiable parameters among StructuralIdentifiability.jl in JULIA and DAISY (Table 2). COMBOS cannot handle the structural stability analyses for <u>Model 4</u>, regardless of whether initial conditions are known or not (Table 2). While increased difficulty in analyzing the structural identifiability of <u>Model 4</u> is not surprising given its increased dimensionality (eight equations), multiple parameters (thirteen) and complex terms of interaction, this model is validated with two data sets (virus titer load and effector CD8^+^ T cells). Our analysis showed that the addition of data for another model variable did not compensate for the size of the model and number of unknown parameters.

Interestingly, we found that all parameters (for all models) are structurally identifiable under known (with the exception of initial effector population, *E*_0_) initial conditions. Given that this is an inoculation study, the assumption of known viral inoculum (set by the experiment) and of known initial target cell population (approximated based on the animal body weight) is not unreasonable. When <u>Models 1-4</u> are used to model natural infection, however, such initial conditions would be unknown and all models would become non-structurally identifiable. Hence, it would be impossible to estimate all parameters even in the presence of unlimited data. A reduction of the unknown parameter space (based on the reported parameter correlations) would be needed before model validation can be attempted.

We next validated <u>Models 1-3</u> with discrete (daily) virus titer data (for the first nine days) and validated Model 4 with discrete (daily) virus titer data (for the first nine days) and discrete (daily) CD8^+^ T cell data (for the first twelve days). Model selection (based on *AIC*_*c*_) favored <u>Model 4</u> as the best model (Table 3). Interestingly, <u>Model 1</u> was the second best model, even though it had the largest 95% error region around the predicted mean virus fit (Fig. 2, grey shaded regions). All models perform worst during the contraction interval (Fig. 2, grey and blue shaded regions), suggesting uncertainty in death rates estimates (for the virus and infected cells).

We used the best parameter estimates obtained through data fitting for <u>Models 1-4</u>, to further investigate their practical identifiablity. Knowing that data used for validation was collected daily and that there was variability among the subjects at each time point, we want to determine wheter there is uncertainty in estimated parameters. When it comes to practical identifiability, several methods are available, such as the *Monte Carlo Simulations* [19–21], the *Fisher Information Matrix* (FIM) or *Correlation Matrix* [16, 22–24], *and Bayesian Methods* [25]. In this study, we used the *Profile Likelihood Method* [26, 38], for two main reasons. First, it allowed us to not just classify the models as practically or non-practically identifiable, but to determine a 95% confidence interval for each practically identifiable parameter. Second, it allowed us to determine the required assumptions needed to improve practical identifiability, while maintaining biological realism (for example, by imposing positivity for all parameters).

We found that none of the models are practically identifiable for the daily empirical data collected in [29] and the parameter range restrictions imposed in Table 1 (see Fig 3A, C, E and Fig 4A). While <u>Model 1</u>, <u>Model 2</u>, and <u>Model 4</u> become practically identifiable if we assume data is collected every fourteen minutes (see Fig 3A, D and Fig 4B), Model 3 does not (see Fig 3F). For this model, we can achieve practical identifiability only when we assume that the viral clearance rate can reach values as high as *c* = 500 per day (corresponding to a life-span for the influenza virus of 2.9 minutes), and that the epithelial cells spend 1*/k* = 1.2 hours in exposed phase before they become productively infected (see Fig. 5). While large influenza clearance rates have been reported before [29], the eclipse phase 1*/k* is assumed to be conserved in a tight interval of 4-6 hours in most respiratory infections [5, 11, 29]. Therefore, this parameter is not practically identifiable from <u>Model 3</u> even when data is collected at high frequency. This is a situation, when a parameter should be removed from the data fitting routine, in order to improve the uncertainty in the estimates of the remaining parameters (see Fig. 6).

**Fig 5.**
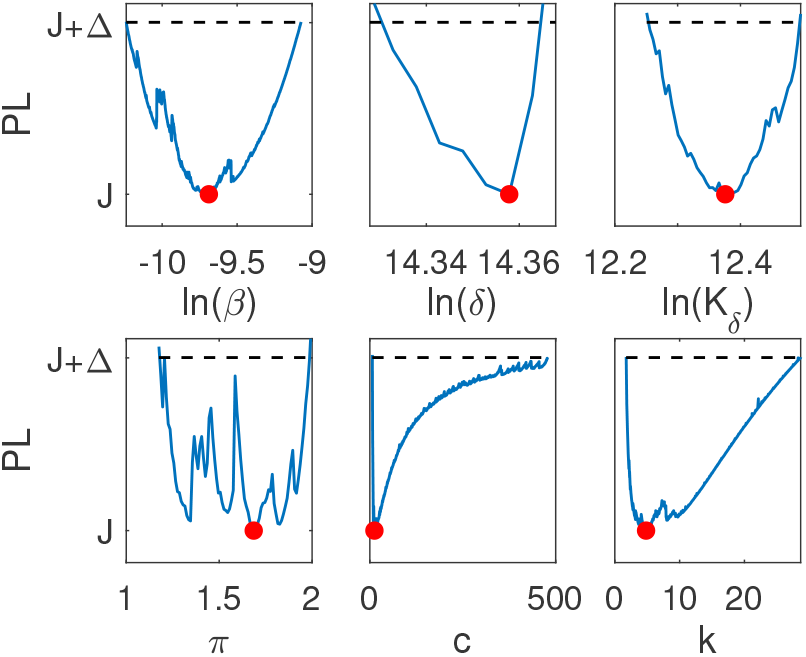
Profile likelihood curves generated using real mice data for <u>Model 3</u> when we relax constraints, such that *c* ∈ [0, 1000] and *k* ∈ [0, 50]. The red circles indicate the best parameter estimate given in Table 3 and the dashed line represents a threshold equivalent to 90% confidence level in the parameter estimate.

**Fig 6.**
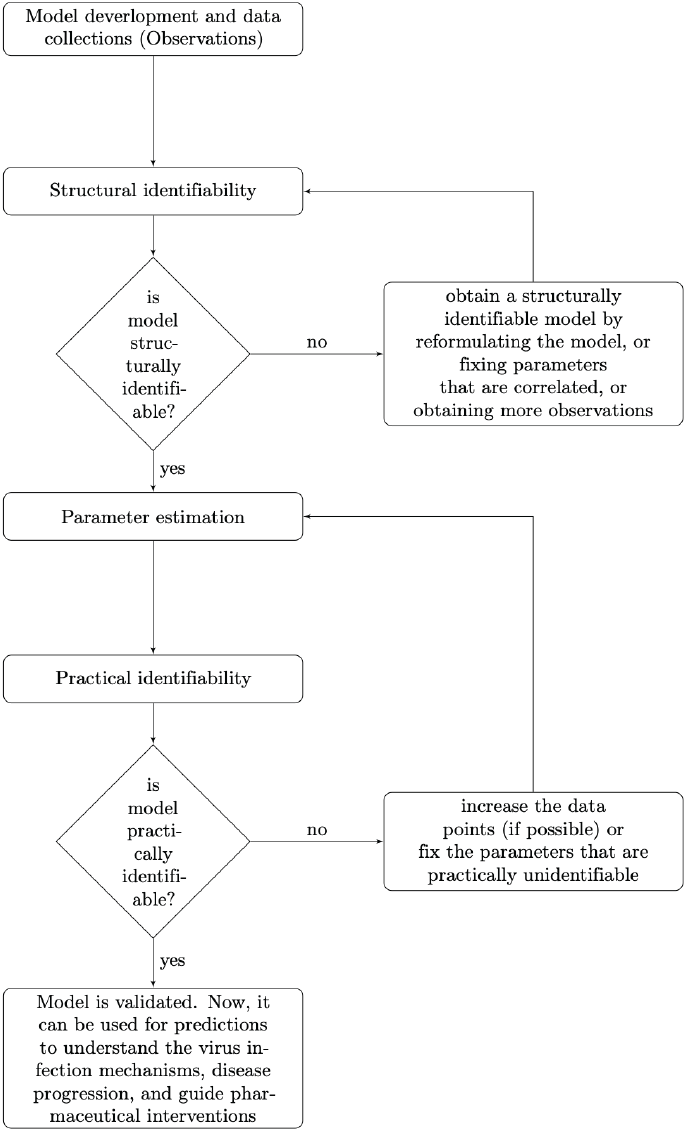
Flow chart of performing identifiability theory to ODE models

Our study has several limitations. First, <u>Model 4</u> was originally expressed as a five order system of delay differential equations. Given the lack of methods that can be used to determine the structural identifiability of delay differential equations, we approximated it with an eight order system of ordinary differential equations. More work is needed to determine whether we maintain (or improve) the practical identifiability results when the delay differential equations system is used in the place of the ordinary differential equations system. Second, we assumed that all model parameters are unknown. It is feasible that the practical identifiability will be improved if certain parameters (such as the eclipse phase) were assumed known. Lastly, all our practical identifiability results come in the context of daily data collection. It would be interesting to see how data collected with random frequency (especially unavailability of data measurements before peak viral load) changes the results.

In conclusion, we investigated the structural and practical identifiability of four nested ordinary differential equation models of influenza A virus infection in mice. We determined the tradeoff between model complexity (defined as combined system dimension, number of unknown parameters, nonlinearity in model interactions), data availability, and our ability to reliably estimate model parameters. We presented solutions for improving model identifiability. While our results dependent on the structure of the models considered and the available data, the methods are generalizable and their use is needed to improve reliability and reproducibility of parameter estimates in other systems of ordinary differential equations applied to discrete biological data.

## Author Contribution Statement

Conceptualization: NT and SMC; analysis: YRL and NHB; funding acquisition SMC and NT; writing - original draft preparation: YRL, NHB, NT and SMC; writing - review and editing: YRL, NHB, NT and SMC.

